# Phylogenetic Analysis of St. Louis Encephalitis Virus within Two Southwestern United State Counties: a case for a bulk introduction event into the southwest United States

**DOI:** 10.1101/2020.06.10.143818

**Authors:** Chase Ridenour, Jill Cocking, Samuel Poidmore, Daryn Erickson, Breezy Brock, Michael Valentine, Steven J Young, Jennifer A Henke, Kim Y Hung, Jeremy Wittie, Heidie M Hornstra O’Neill, Krystal Sheridan, Heather Centner, Darrin Lemmer, Viacheslav Fofanov, Kirk Smith, James Will, John Townsend, Paul S. Keim, David M. Engelthaler, Crystal M Hepp

**Affiliations:** School of Informatics, Computing, and Cyber Systems, Northern Arizona University, Flagstaff, Arizona, United States of America; The Pathogen and Microbiome Institute, Northern Arizona University, Flagstaff, Arizona, United States of America; Translational Genomics Research Institute, Flagstaff, Arizona, United States of America; Maricopa County Environmental Services Department Vector Control Division, Phoenix, Arizona, United States of America; Coachella Valley Mosquito and Vector Control District, Indio, CA

## Abstract

St. Louis Encephalitis Virus (SLEV) has been seasonally detected within the *Culex spp*. populations within Maricopa County, Arizona and Coachella Valley, California since an outbreak in Maricopa County in 2015. Previous work revealed that the outbreak was caused by an importation of SLEV genotype III, which had only been detected within Argentina in prior years. However, little is known about when the importation occurred or the population dynamics since its arrival into the southwestern United States. In this study, we wanted to determine if the annual detection of SLEV in Maricopa and Riverside counties is due to enzootic cycling or new importations. To address this question, we analyzed 143 SLEV genomes (138 sequenced as part of this study) using the Bayesian phylogenetic analysis software, BEAST, to estimate the date of arrival into the American Southwest and characterize the underlying population structure of SLEV. Phylogenetic clustering showed that SLEV variants circulating in Arizona and California form two distinct populations with little evidence of transmission among the two populations since the onset of the outbreak. Interestingly, the SLEV variants in Coachella Valley appear to be annually imported from a nearby source, whereas the Arizona population is locally sourced each year. Finally, the earliest representatives of SLEV genotype III in the southwestern US formed a polytomy that includes both California and Arizona samples. We propose that the initial outbreak could have resulted from an introductory population of SLEV, perhaps in one or more bird flocks migrating north in 2013, rather than a single variant introduced by one bird.

## Introduction

St. Louis Encephalitis Virus (SLEV) is the causative agent of the disease St. Louis Encephalitis (SLE). Transmission of SLEV is enzootically cycled between *Culex spp*. mosquito vectors and amplified in numerous bird hosts [1][2]. While SLEV can infect humans, it does not achieve high enough levels of viremia to be further transmitted. SLE symptoms include headache, fever, nuchal rigidity, disorientation, and tremor, and is often confused with the flu. However, 80% of clinical cases result in encephalitis, of which 5-20% are fatal [1]. SLEV has been circulating in the Americas for nearly 300 years [1]; yet, it was not discovered until 1933, after a viral outbreak in St. Louis, Missouri resulted in 1095 reported cases, including 201 fatalities [3]. Since 1933, over 50 SLEV outbreaks have occurred in the United States and southern Canada, resulting in ~10,000 reported encephalitis cases and more than 1,000 fatalities [2,3].

SLEV’s incidence and medical prominence within the United States was displaced with the importation of West Nile Virus (WNV) in 1999 [4,5]. WNV quickly spread throughout the United States, becoming established in the US by 2004 [6]. Concurrent with the geographic radiation of WNV, SLEV cases caused by genotypes I and II rapidly decreased [4][7]. With both SLEV and WNV being heterologous flaviviruses, it is well-understood they impart cross-immunity between shared hosts [8,9]. Previous work revealed that prior infection with WNV can cause complete immunity to a secondary SLEV genotype II or V infection, whereas a primary SLEV infection reduced the viral load of WNV by a thousand fold [9–12]. The leading hypothesis for the drastic decrease in cases of SLEV is that there are cross-immunity effects between WNV and previously circulating genotypes of SLEV [4].

In 2015, a SLEV outbreak occurred in Maricopa County, Arizona where 23 individuals were infected, 19 of whom developed encephalitis, resulting in two fatalities [12,13]. These were the first cases of SLEV in Arizona in 10 years. Moreover, this was the first outbreak in the United States since the 2001 Louisiana outbreak that resulted in four deaths [14]. The SLEV strain isolated from the 2015 epidemic shared sequence homology with Argentinian strains, placing it within Genotype III, the same genotype that caused the first epidemic in South America [13][15]. During 2005, Cordoba City, Argentina had 47 probable human cases of SLEV which resulted in 45 hospitalizations and nine deaths [15]. This is not the first time a South American genotype was observed circulating within the United States [16][17]. An SLEV strain collected in Florida in 2006 was determined to be genotype V which was previously only observed within South American[16][17]. The current working hypothesis is the South American genotypes have been introduced via migratory birds.

To date, it is unknown whether the seasonal observations of SLEV within Maricopa County are due to the virus overwintering locally or from annual importations from surrounding regions [7,18]. Therefore, acquiring a thorough understanding of the molecular history and epidemiology of SLEV is essential information for health agencies to assess the potential risk of SLEV to the local populations [12]. The overarching goals of our study were to determine the time of entry of SLEV into the Southwest US and determine the spatial-temporal circulatory trajectories within Maricopa County, AZ and Riverside County, CA. Our central hypotheses were that a single SLEV migration event occurred several years prior to the 2015 outbreak, and SLEV has become endemic to both counties’ mosquito populations [12,19].

## Material and Methods

### Sample Collection

The Maricopa County Environmental Services Vector Control Division (MCESVCD) and Coachella Valley Mosquito and Vector Control District (CVMVCD) conduct regular mosquito trapping and abatement throughout their respective districts. MCESVCD mosquito surveillance program places ~800 CO2 traps throughout the Phoenix Metropolitan area. Each trap is placed within its designated square mile area (640 acres) for a 12-hour collection period once weekly.

The CVMVCD mosquito surveillance program splits Coachella Valley into two regions: eastern valley and western valley. The eastern valley has 56 CO2 traps set every two weeks. The western valley has 53 gravid traps and 53 CO2 traps weekly. After collection, mosquitoes are sorted by sex, with males being discarded, while the females are pooled by species with a maximum of 5 pools per trap and 50 females per pool. Resulting pools are tested for WNV and SLEV by MCESVCD and CVMVCD, following the protocol described by Lanciotti et al. [19]. SLEV-positive pools are stored in −80°C freezers until they are either same-day transported to Northern Arizona University on dry ice (MCESVCD) or shipped using FEDEX ground transportation while being preserved in DNA/RNA ShieldTM 2X Concentrate. Metadata supplied with each of the samples includes the GPS coordinates of the mosquito traps, date of collection, total number of mosquitoes captured, and mosquito species. For this study, we selected a total of 138 positive mosquito pools for whole genome sequencing. The MCESVCD set included 28 positives in 2015, 24 in 2017, 14 in 2018 and 41 in 2019. The CVMVCD set was composed of 16 positives in 2017 and 15 samples from 2018. Samples were selected to capture the spatial and temporal diversity of SLEV within Maricopa County and Coachella Valley.

### Sample Processing, SLEV Tiled amplicon sequencing

The methods used to prepare the SLEV samples for transport, RNA extraction, and reverse transcription followed the protocol previously described for WNV by Hepp et al. [19,20]. The multiplex PCR primers were designed using the software package Primal Scheme [21], where the 42 primer pairs were based on a genome from Kern Valley, California (KY825743.1) with an average primer pair product of 400bp. For each sample, a multiplex PCR reaction was performed for each pool individually. The PCR reaction used 12.5μL of KAPA 2G Fast Multiplex Mix (2X) (Kapa Biosystems, Wilmington, MA), with a final primer concentration of 0.2 μM, 2.5 μL of cDNA,in a total reaction volume of 25 μL. The thermocycler settings used: 3 minutes of denaturation at 95°C f, 30cycles of 98°C for 15 seconds, 60°C for 30 seconds, 72°C for 1 minute, and a final extension of 72°C for 1 minute. The PCR product was cleaned using 1X Agencourt AMPure XP beads (Beckman Coulter, Indianapolis, IN). A second PCR using universal tail-specific primers was performed to add the Illumina specific indexes [22]. The reagents for the reaction were 12.5 μL of 2X Kapa HiFi HotStart Ready Mix (Kapa Biosystems), 400 nM of each forward and reverse indexed primer, and 2 or 4 μl of the cleaned amplified SLEV product. The thermocycler protocol is as follows: 98°C for 2 minutes, 6 cycles of 98°C for 30 seconds, 60°C for 20 seconds, 72° for 30 seconds, and a final extension at 72°C for 5 minutes. The DNA for the samples in each pool was quantified using the Kapa Library Quantification kit (Kapa Biosystems). The samples were then pooled to achieve an equal concentration of each sample. Sequencing was conducted on the Illumina Miseq sequencing platform, using a v3 600 cycle kit.

### Post-sequencing Data Processing

To generate consensus sequences needed for phylogenetic analysis, sequencing reads were first trimmed using Amplicon Sequencing Analysis Pipeline 0.9 (ASAP) (https://github.com/TGenNorth/ASAP). ASAP first trimmed the reads of adapter and primer sequences using BBDuk, a tool integrated into the BBMap package (https://sourceforge.net/projects/bbmap/). Resulting paired-trimmed reads were aligned to the reference genome, FJ753286.2, using Bowtie2 [23], and alignments were indexed using Samtools 1.4.1 [24]. Consensus sequences were generated using the program iVar 1.0 [24,25]. The criterion for base calling was a minimum of 10x coverage and a majority base proportion of 0.80. Base calls were coded according to the International Union of Pure and Applied Chemistry (IUPAC) nucleotide codes. Additionally, a read pileup was produced using the Integrated Genome Viewer(IGV) 2.4.16 command line tool count [26].

### Maximum Likelihood Analysis

To determine if SLEV strains circulating within Maricopa County and Coachella Valley were genotype III, we conducted a Maximum-Likelihood phylogenetic analysis using the 44 publicly available whole genome sequences from the National Center for Biotechnology Information (NCBI) and the 138 genomes sequenced within our lab. Sequences were aligned using MUSCLE [27] followed by substitution model testing using IQ-TREE. The F81 substitution model with empirically predicted base frequencies was found to be the best fit model based on the Bayesian information criterion. The maximum-likelihood tree was constructed using IQ-TREE with 1000 bootstraps iterations [28][28,29][30]. The tree was rooted using the two strains of SLEV (JQ957869.1 and JQ957870.1) detected in Columbia in 2008 as described by Hoyos-Lopez et al. [31].

### Bayesian Phylogenetic Analysis

The temporal signal of the 143 SLEV genomes was determined by first generating a maximum likelihood (ML) phylogeny with IQ-TREE using the F81 substitution model [32] with empirically generated base variation, which was determined by IQ-TREE’s model selection. Then, a root-to-tip genetic divergence and time of sampling regression was performed in TempEst v1.5.1 [33].

To estimate the time of entry and population structure of SLEV in the American Southwest, a Bayesian phylogenetic analysis was conducted using the BEAST v1.10.4 software package [33,34]. The specified substitution model, as determined by IQ-TREE previous, is the F81 nucleotide substitution model with empirically derived base frequencies. The best fitting molecular clock and demographic model were determined by marginal likelihood comparison using path-sampling and stepping-stone sampling [35][35,36], see S1 Table. The Bayesian Skyride model was found to be best fitting in combination with a relaxed molecular clock. The final model was run on four independent Markov chains where each chain ran for 100 million Markov chain Monte Carlo steps and the state was sampled every 10,000 steps. Convergence was assessed using Tracer v1.7 [33,34,37]. States were combined using LogCombiner, discarding the first 10% as burn-in (10,000,000 generations per chain), and then resampling every 30,000 generations. Trees were summarized using TreeAnnotator [8], producing a maximum clade credibility (MCC) tree that was visualized using the R package GGtree [38][39][40]. The branches of the tree were collapsed into polytomies if the posterior support was below 0.80 using a custom R script https://github.com/ChaseR34.

## Results

### Genetic Relatedness of 2015 to historical

To understand the genetic relatedness of the 2015 outbreak, a maximum likelihood phylogenetic analysis was conducted using 184 SLEV genomes (138 of the genomes were produced by our lab and 44 previously sequenced SLEV genomes (Fig. 2). All Maricopa County (blue) and Coachella Valley (orange) samples collected during or after the 2015 outbreak formed a monophylectic clade, referred to as the outbreak clade, nested under the 2005 Genotype III Argentinian samples (Figs. 2a and c). Furthermore, the 2015 outbreak clade is distinct from the previously endemic United States genotypes I and II (purple, Figs. 2a and b). Collectively, this analysis provides further evidence that the introduction of Genotype III into the southwest United States was due to a single migration event into the southwestern United States, corroborating conclusions made by White et al. [13] and Diaz et al. [7].

### BEAST Phylogenetic Analysis

#### Date of entry and bulk migration event

The first line of inquiry was to estimate the timing and location for the initial SLEV genotype III migration into the southwest United States. To estimate these values, we conducted a Bayesian phylogenetic analysis using the software BEAST. The most recent common ancestor of all United States-based Genotype III SLEV strains is well-supported, and we estimate the migration event into the United States to have occurred between September 2011 and May 2014 (median 2013.5, 95% HPD CI:2011.71-2014.34). The phylogeny further supports three distinct and well-supported clades (posterior probability > 0.80): hereafter referred to as Clade 1, Clade 2, and Clade 3 (Fig 3). However, the branch support for several interior branches is poor (posterior value < 0.80, Figure S1) resulting in the forced polytomies observed in Figure 1.

**Figure 1:**
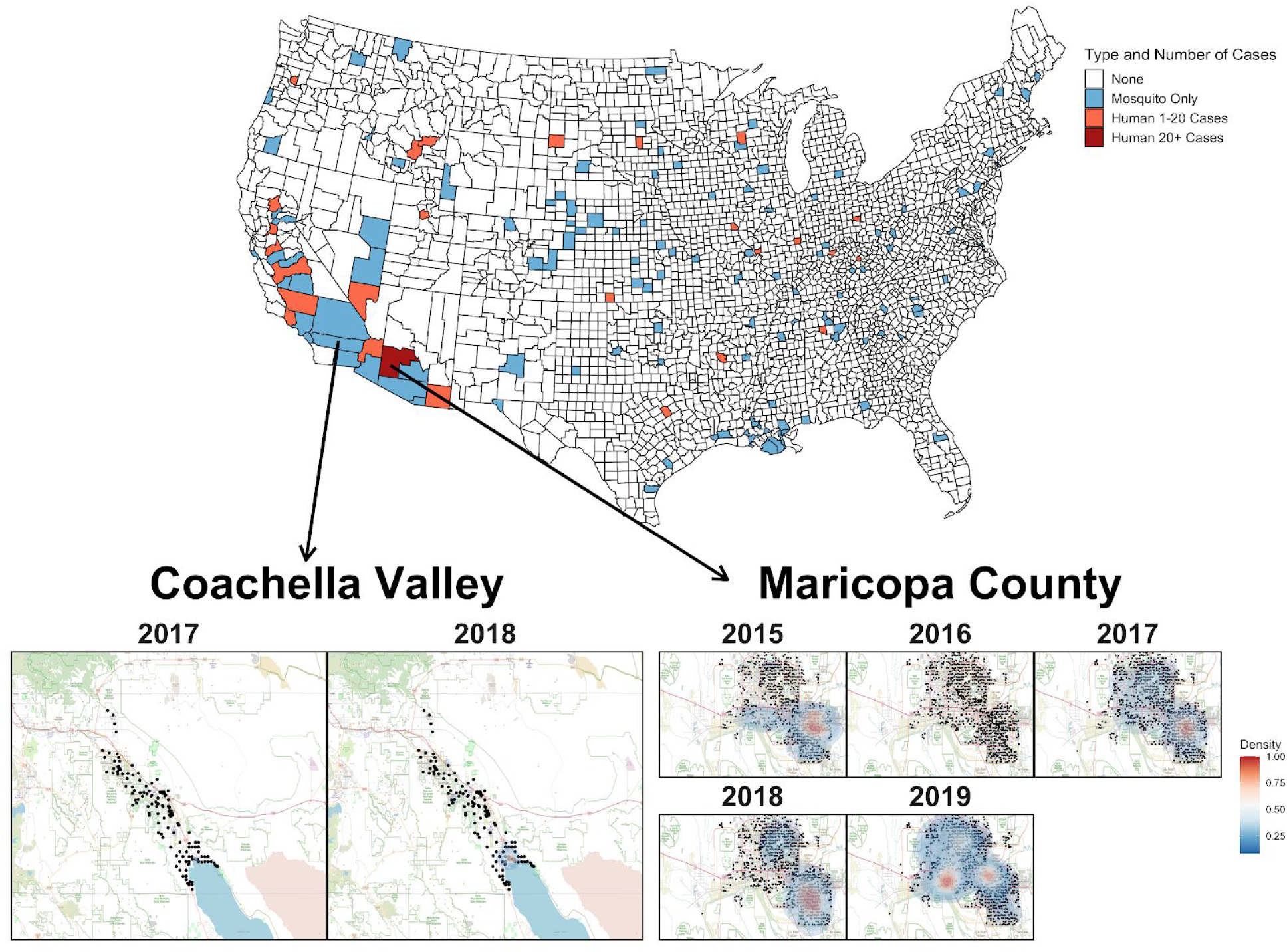
The US map colors counties where SLEV positive mosquitoes or human cases were observed between 2015 and 2019. The blue counties indicate SLEV was detected within mosquito populations only. The red counties indicate human cases and the dark red county is Maricopa County which is the only county with over 20 human cases. Heat map of densities of positive mosquito traps found within Coachella Valley between 2015-2018 and Maricopa County between 2015-2019 and. The black points represent the individual mosquito trap locations. A two-dimensional kernel density estimation with axis-aligned bivariate normal kernel was used to estimate the density of positive mosquito traps within each year. Red indicates a higher density of positive traps.

**Figure 2:**
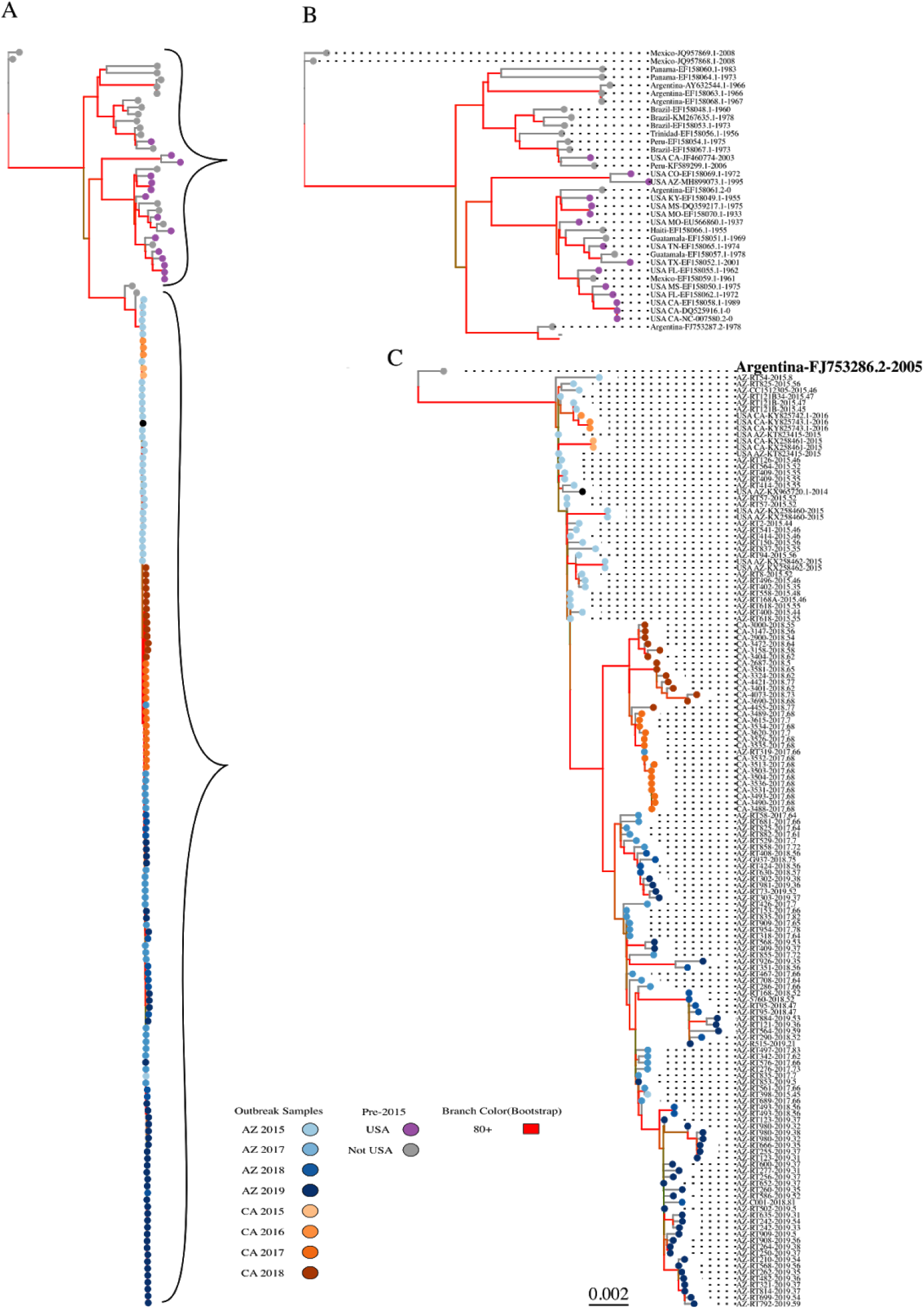
A) The Maximum Likelihood Tree constructed from 184 SLEV genomes (44 public genomes and 140 genomes produced by our lab). The blue and orange tips are the post-2015 samples, purple are the local USA genotypes I and II, and grey are the Central and South American sourced genomes. B) The dashed arrow points to a closeup of the clade consisting of the USA genotype I and II and Central and South American strains. C) Closeup of the 2015 outbreak clade. Tips are labeled with country or state of origin, NCBI accession number for publicly available genomes or sample ID for unpublished sequences, and the date of sampling to the nearest hundredth of a year. All the post-2015 samples form a monophyletic clade nested under the Argentina_FJ753286.2_2005 sample indicating all post 2015 outbreak samples are of genotype III.

**Figure 3:**
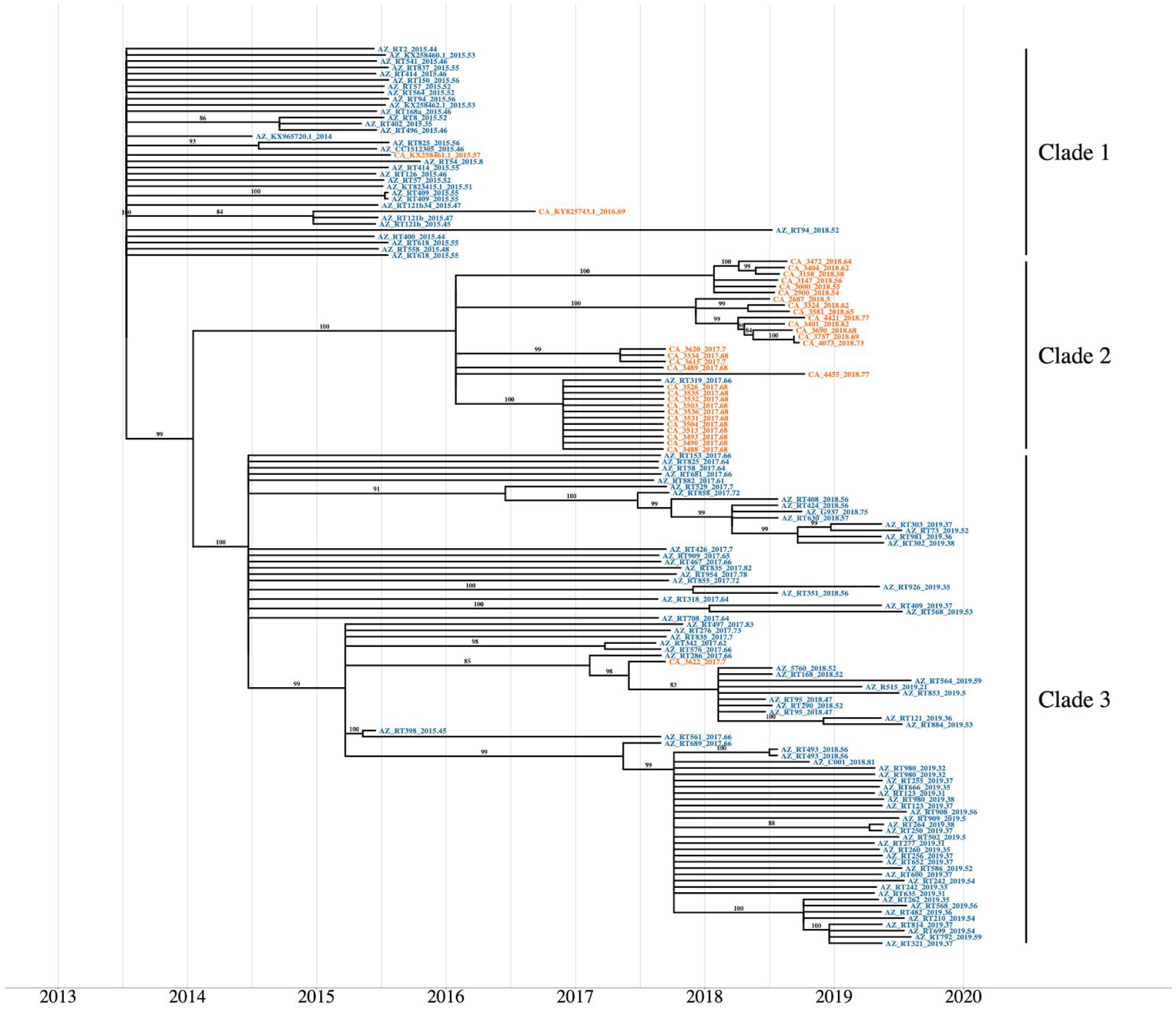
The maximum clade credibility phylogenetic tree reconstructed using 144 genotype III SLEV genomes from Arizona and California. The tip colors distinguish the sampling location of each sample, Arizona (Blue) and California (Orange). The posterior values for branches above 80 are written above their corresponding branch. Branches with posterior values below 80 were collapsed into polytomies. After correcting for low confidence branches, the phylogeny clusters into three distinct clades. Clade 1 consisting of all the 2015-2016 California and 2015 Arizona samples. Clade 2 consists of the 2017-2018 California samples. Clade 3 contains the 2017-2019 Arizona samples.

Clade 1 is the most ancestral clade, therefore, the inference concerning entry location and timing will be drawn using Clade1’s topology. The confidence estimates on clusters within Clade 1 are too low (<0.5) to make conclusions regarding which southwestern location SLEV genotype III first entered, which can be seen by the majority of the samples forming polytomies (Figs. 1 and S1). The observance of polytomies is indicative of samples within the clade constituting a diverse genetic population. Moreover, Clade1 contains genomes from both Arizona and California which do not form any clear nesting, therefore, we are unable to distinguish if SLEV arrived into either California or Arizona first. However, the lack of clear structure within Clade 1, leads us to consider the possibility that the introduction of SLEV into the southwestern United States was a bulk migration event over a short period of time, e.g. a flock of birds with multiple infected individuals, which would explain the lack of a robust signal indicating a single entrance into one location over another.

#### Characterizing Source Location of Seasonal SLEV Populations

Our second line of inquiry was to determine if the seasonal occurrences of SLEV within Maricopa County, Arizona, and Coachella Valley, California are sourced from the endemically circulating variants or are newly introduced via yearly migration events. These migration events will appear as specific branching patterns within the major clades. Since, we are interested in estimating migration patterns for current populations, only the two clades containing extant lineages, Clade II composed primarily of Coachella Valley samples, and Clade III containing Maricopa County samples, will be considered. The geographic clustering allows us to draw two conclusions: first, that migration events between Coachella Valley and Maricopa County have been rare, and second, the inference about the two populations can be done independently. Therefore, the two clades are interpreted separately below.

Clade II contains the Coachella Valley, California samples from 2017 to 2018. The internal branch topology of Clade II supports two monophyletic clusters, distinguished by year of collection. This topology is indicative of two possible scenarios for the seasonal appearance of SLEV. The first scenario is that the population of SLEV in 2018 was seeded by cryptically circulating SLEV from the previous year. The second, more plausible scenario, is that a closely related, and perhaps closely situated, population of SLEV migrated into Coachella Valley each year.

Clade III is composed of the 2017, 2018, and 2019 Arizona samples, and a single CA sample. There are two plausible patterns to the seasonal establishment of SLEV within Maricopa County. First, if SLEV was sourced from nearby regions, rather than internally, we would expect to observe multiple monophyletic clades that cluster by year, mirroring the Clade II topology. The second possibility is a temporally paraphyletic clade that includes the 2018 and 2019 AZ samples nesting within the 2017 AZ samples. The phylogeny clearly shows Clade III is temporally paraphyletic with the 2018 and 2019 samples being nested within 2017 samples across multiple internal clades. Therefore, our results indicate SLEV has become endemic in Maricopa County and the seasonal emergence of SLEV has been annually re-seeded by multiple local populations within Maricopa County.

## Discussion

The focus of our study was to better understand the dynamics of St. Louis Encephalitis Virus circulation in the southwest United States since the 2015 outbreak. We sequenced a total of 138 genomes from Coachella Valley, California and Maricopa County, Arizona from 2015 through 2019. Using Bayesian phylogenetic techniques, we considered the following: 1) when the first introduction of genotype III into the southwest United States occurred, 2) the number of distinct introductions that have occurred, 3) whether contemporary variants in Maricopa County or Coachella Valley have become endemic, and 4) the amount of time the endemic variants have been establishment.

Our study revealed that the migration of SLEV into the southwest United States occurred between September 2011 and May 2014 (median 2013.5, 95% HPD CI:2011.71-2014.34), but went undetected until the 2015 outbreak. Our working hypothesis is SLEV was circulating at low levels within the avian populations of the southwestern United States escaping detection. Since the 2015 outbreak, SLEV has been reliably circulating within the Southwestern United States. However, an interesting caveat was in 2016 where Maricopa County, the epicenter of the 2015 outbreak, reported zero human cases or positive mosquito traps. We are still uncertain the exact cause of this population crash of SLEV in Maricopa County. Then in 2017, SLEV was detected at high levels and has sustainably been detected ever since. The phylogeny in Fig 1 strongly insinuates the reemergence of SLEV in Maricopa County was seeded by low-level circulating strains of SLEV. The fact that the Clade II (California) and Clade III (Arizona) are isolated populations with very little migration between the two populations indicates there is very low likelihood that a migration event from California seeded the 2017 reemergence in Maricopa County. Furthermore, only a single AZ sample, AZ_RT319_2017.66, and a single CA sample, CA_3622_2017.7, do not cluster within their expected clades; neither have extant lineages. If a migration event into Maricopa County was what seeded the reemergence in 2017 was from California we would expect to see extant lineages deriving from a common California ancestor which is not observed from the phylogeny. Therefore, We are very confident that the 2017 reemergence in Maricopa County was due to local low-level circulating strains of SLEV. Since 2017, SLEV has been locally circulating within Maricopa County.

Unlike Maricopa County, the Coachella Valley sample’s seasonal circulation is unclear whether it is seeded by the immigration of SLEV from a nearby endemic population or if there is endemic low level circulation within Coachella Valley. The Salton Sea is on the flight path of many migratory birds and is an important water source for local bird populations. This fact along with the formation of monophyletic clades for 2017 and 2018 form within Clade II leads us to tentatively conclude that transient strains of SLEV are responsible for the seasonal circulation within Coachella Valley. However, samples from other regions of California are needed to conclusively determine the origin of Coachella Valley’s seasonal SLEV populations.

Overall, genotype III SLEV has been the majority genotype infecting humans since its introduction in 2015. Between 2015 and September 2019, 50 of the 59 human cases of SLEV reported nationwide to the Center for Disease Control (CDC) were genotype III and occurred within Arizona and California. Furthermore, in 2019, the CDC reported the highest SLEV environmental load ever in the southwest United States. In Maricopa County, the SLEV record environmental load is occurring alongside WNV having a near record year. Coachella Valley had a record breaking year as well, with 513 WNV and 105 SLEV samples. These levels of co-circulation between SLEV and WNV have not been previously observed in the United States. Therefore, SLEV appears to be a consistent health risk in the southwestern US.

