## Supplemental Info for "Phylogenetic Analysis of St. Louis Encephalitis Virus within Two Southwestern United State Counties: a case for a bulk introduction event into the southwest United States"

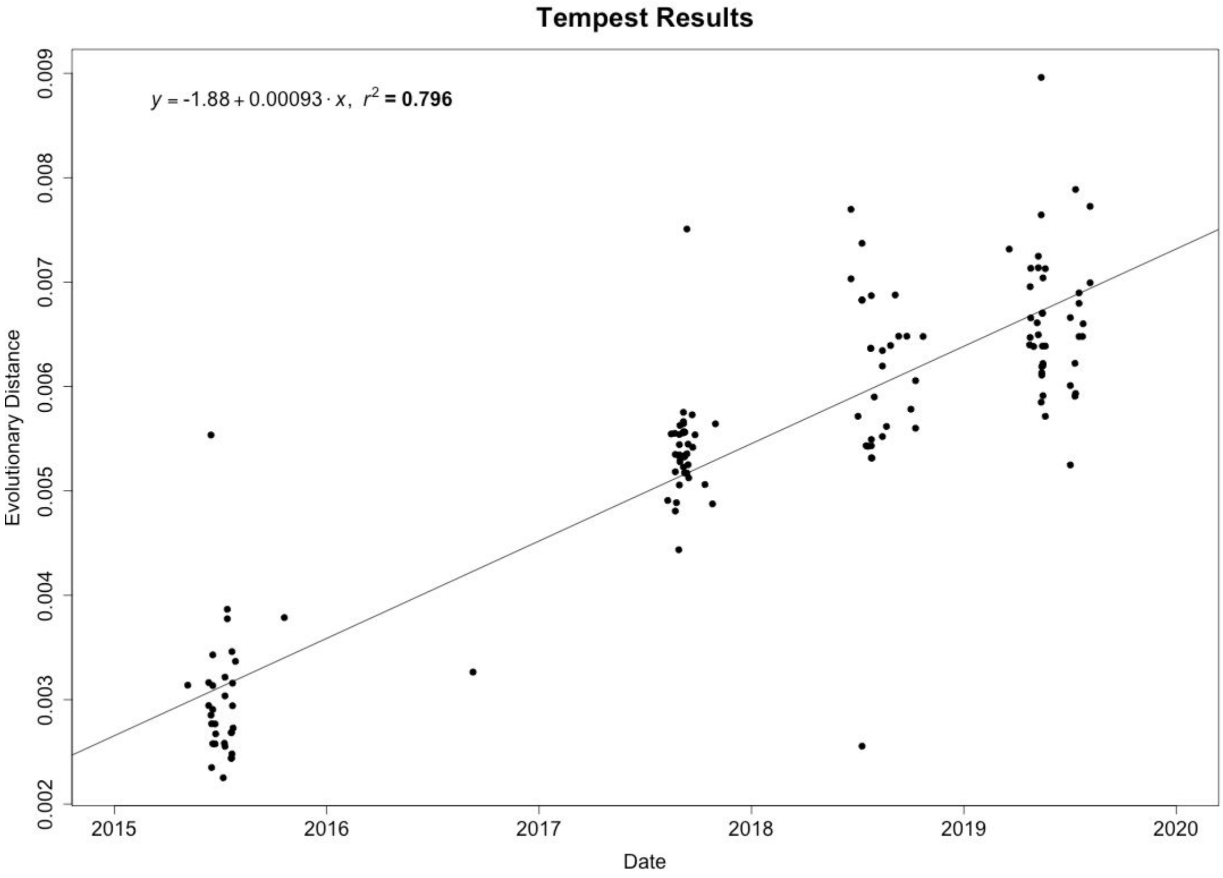


*Figure S1: Linear regression of genetic distance versus time for all 146 genomes used to determine if a molecular signal was detectable*

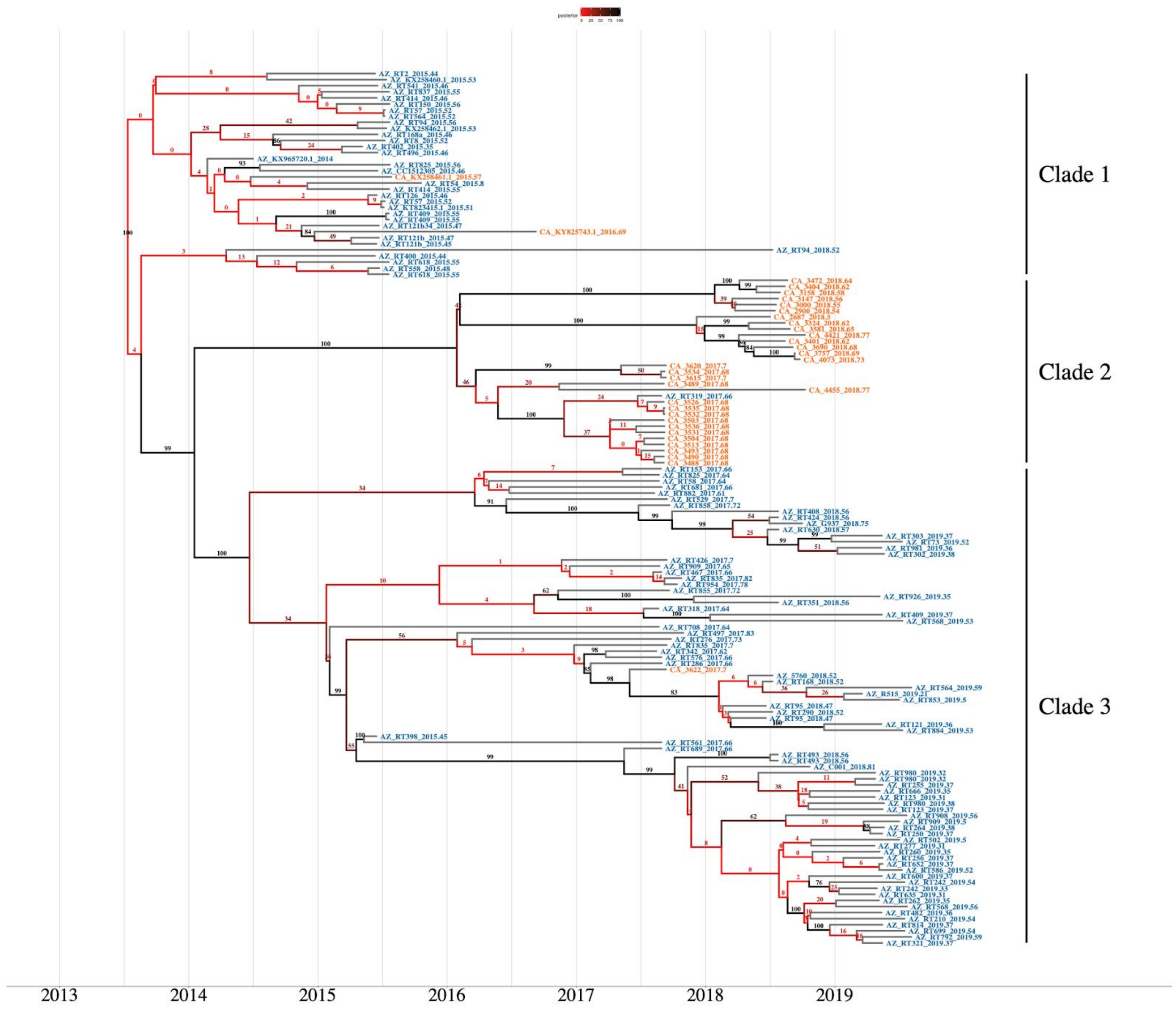


*Figure S3:* The maximum clade credibility phylogenetic tree reconstructed using 146 genotype III SLEV genomes from Arizona (blue) and California (orange). This is the same tree as Figure 1 but prior to collapsing low-confidence branches into polytomies. The red branches indicate low posterior probability and the black branches indicate high posterior probability. The exact posterior value is given by the number on the branch.

Table S2: BEAST Model Testing using generalized stepping-stone sampling

| | Strict Molecular Clock | Relaxed Molecular Clock Lognormal |
|---|---|---|
| Constant Population | -20648.803 | -20555.859 |
| Exponential Population | -20656.096 | -20558.3066 |
| SkyGrid | -20648.674 | -20554.049 |
| Skyride | -20688.523 | **-20547.215** |

Table S2: SLEV Multiplex PCR Primer Pairs

| Primer.Name | Sequence | Len | Tm | GC | Start | End |
|---|---|---|---|---|---|---|
| SLEV_1_LEFT | GGTGAGCGGAGAGGAAACAGAT | 22 | 62.04 | 54.54 | 13 | 35 |
| SLEV_1_RIGHT | TCCCTCCTCTCTTCTTGCTTGG | 22 | 61.42 | 54.54 | 414 | 392 |
| SLEV_2_LEFT | AGCCATCCTGACATTCTTCCGA | 22 | 61.47 | 50 | 241 | 263 |
| SLEV_2_RIGHT | CACCAACAGTCAATGTCCTCGG | 22 | 61.43 | 54.54 | 667 | 645 |
| SLEV_3_LEFT | CTAGTGCCAACGGAGCAAACAC | 22 | 62.25 | 54.54 | 537 | 559 |
| SLEV_3_RIGHT | ACCACTCTCTGTGTGTTGTTGC | 22 | 61.25 | 50 | 922 | 900 |
| SLEV_4_LEFT | GGACACCGTGAAAACCACCAAA | 22 | 61.78 | 50 | 796 | 818 |
| SLEV_4_RIGHT | CGTGTCCAAGGTTGCTTCGTAA | 22 | 61.36 | 50 | 1163 | 1141 |
| SLEV_5_LEFT | TACTTGAAGGGGGAAGCTGTGT | 22 | 61.29 | 50 | 1032 | 1054 |
| SLEV_5_RIGHT | CGTAGAGTCCGTTGAACCATGC | 22 | 61.54 | 54.54 | 1412 | 1390 |
| SLEV_6_LEFT | GCCTGTTTGGAAAAGGGAGCAT | 22 | 61.67 | 50 | 1278 | 1300 |
| SLEV_6_RIGHT | GCTCGTCCATGGAAGGTTCAAG | 22 | 61.50 | 54.54 | 1643 | 1621 |
| SLEV_7_LEFT | GGAACAGTTACCATTGATTGTGAAGC | 26 | 61.06 | 42.31 | 1511 | 1537 |
| SLEV_7_RIGHT | CAGTTCCACAATCACTGTCCCG | 22 | 61.43 | 54.54 | 1943 | 1921 |
| SLEV_8_LEFT | ATGCAGAGCTAAGCTTGACAAGG | 23 | 61.18 | 47.82 | 1822 | 1845 |
| SLEV_8_RIGHT | AAGCCTTACCAATGCTGCTTCC | 22 | 61.46 | 50 | 2184 | 2162 |
| SLEV_9_LEFT | GGGAGCGAACAACAAGGTCATG | 22 | 62.02 | 54.54 | 2056 | 2078 |
| SLEV_9_RIGHT | AGCCAGTAGAGTCAGCGAGATG | 22 | 61.59 | 54.54 | 2423 | 2401 |
| SLEV_10_LEFT | CCACCAAGTTTTCGGAGGAG | 22 | 61.13 | 54.54 | 2285 | 2307 |
| SLEV_10_RIGHT | CGTTGATCGGATGCCACAGATG | 22 | 61.93 | 54.54 | 2645 | 2623 |
| SLEV_11_LEFT | GGAGGAGGCATCTTCGTGTACA | 22 | 61.78 | 54.54 | 2510 | 2532 |
| SLEV_11_RIGHT | TCCATGCTCTGTTTGCTGTTGG | 22 | 61.64 | 50 | 2916 | 2894 |
| SLEV_12_LEFT | GCTGGAGGATGAATTGGACTACG | 23 | 61.04 | 52.17 | 2782 | 2805 |
| SLEV_12_RIGHT | CACCGGGATGATCATTTCGCTT | 22 | 61.32 | 50 | 3200 | 3178 |
| SLEV_13_LEFT | AGAGCAAAAAGAATGAGACATGGCA | 25 | 61.50 | 40 | 3072 | 3097 |
| SLEV_13_RIGHT | GGCCGGATTTCCATTCCATACC | 22 | 61.39 | 54.54 | 3475 | 3453 |
| SLEV_14_LEFT | AACATTGTGGAAACAGGGGAGC | 22 | 61.34 | 50 | 3330 | 3352 |
| SLEV_14_RIGHT | CCTCCAGTGTTCATTTCCGCAA | 22 | 61.39 | 50 | 3736 | 3714 |
| SLEV_15_LEFT | AAGCTGACCCTGACCTCACTAG | 22 | 61.14 | 54.54 | 3611 | 3633 |
| SLEV_15_RIGHT | GCCATCCTCATTCCTGGAGTCA | 22 | 61.82 | 54.54 | 4015 | 3993 |
| SLEV_16_LEFT | TGAAGCTTGAGGTCCTTCCGAT | 22 | 61.42 | 50 | 3882 | 3904 |
| SLEV_16_RIGHT | GCTATTGCAAAAGGGACCACCA | 22 | 61.40 | 50 | 4309 | 4287 |
| SLEV10918_15_LEFT | AAGTGAGGTCTTGACCGGAGTC | 22 | 61.65 | 54.54 | 4149 | 4171 |
| SLEV_17_RIGHT | AATGGAATCGTGCACTCAAGCC | 22 | 61.77 | 50 | 4539 | 4517 |
| SLEV_18_LEFT | CCCAGGCTTGATGTTGACCTTG | 22 | 61.71 | 54.54 | 4418 | 4440 |
| SLEV_18_RIGHT | CGGACATCTCCTGCATACGGAT | 22 | 61.97 | 54.54 | 4813 | 4791 |
| SLEV10918_17_LEFT | GGAGAAGGGAGACTAGATCCGT | 22 | 60.15 | 54.54 | 4711 | 4733 |
| SLEV_19_RIGHT | CTTTCCCCTTGGATGATGCCAC | 22 | 61.53 | 54.54 | 5104 | 5082 |
| SLEV_20_LEFT | CCGTGACGCTTGATTTCCCAAA | 22 | 61.96 | 50 | 4968 | 4990 |
| SLEV_20_RIGHT | GCCCTGATGTTCATTCCTCACG | 22 | 61.57 | 54.54 | 5357 | 5335 |
| SLEV10918_19_LEFT | ACACCAGCCGTGAAGAATGAAC | 22 | 61.39 | 50 | 5263 | 5285 |
| SLEV_21_RIGHT | CCTGGGCCTCAACATCCAGTAT | 22 | 61.82 | 54.54 | 5613 | 5591 |
| SLEV_22_LEFT | GCATTGCTGCTCGTGGGTATAT | 22 | 61.13 | 50 | 5475 | 5497 |
| SLEV_22_RIGHT | TGACTGGTTTCACACACTTGCG | 22 | 61.82 | 50 | 5895 | 5873 |
| SLEV_23_LEFT | GGAAAAGCTTTGACACAGAATACCCT | 26 | 61.68 | 42.31 | 5757 | 5783 |
| SLEV_23_RIGHT | ATCGAAACTCCCCATCCATGGT | 22 | 61.50 | 50 | 6168 | 6146 |
| SLEV_24_LEFT | CATGATCTGGCCAACTGGACTG | 22 | 61.25 | 54.54 | 6041 | 6063 |
| SLEV_24_RIGHT | ATGACTTGAGCGCTTGGTAGTC | 22 | 60.68 | 50 | 6423 | 6401 |
| SLEV10918_23_LEFT | GACTACCAAGCGCTCAAGTC | 22 | 61.49 | 54.54 | 6335 | 6357 |
| SLEV_25_RIGHT | CATCACAAATGCTCCCAGTCCC | 22 | 61.53 | 54.54 | 6710 | 6688 |
| SLEV10918_24_LEFT | GCTGGGAGCATTTGTGATGACT | 22 | 61.20 | 50 | 6627 | 6649 |
| SLEV10918_24_RIGHT | GGATGCGATGGCTGTTAGTGAG | 22 | 61.36 | 54.54 | 7060 | 7038 |
| SLEV_27_LEFT | ACTGGCTGTCTTCTTGATATGCA | 23 | 60.04 | 43.47 | 6844 | 6867 |
| SLEV_27_RIGHT | TTGGCAGGGTCATCTGATTCCA | 22 | 61.70 | 50 | 7233 | 7211 |
| SLEV_28_LEFT | TAACAGCCATTGCATCCCAAGC | 22 | 61.78 | 50 | 7110 | 7132 |
| SLEV_28_RIGHT | CTGCTGACCCTAAGACCCCAAA | 22 | 61.95 | 54.54 | 7518 | 7496 |
| SLEV_29_LEFT | GCGACCCCAATGACAGAGAAGAA | 23 | 62.74 | 52.17 | 7397 | 7420 |
| SLEV_29_RIGHT | GGGCTCTGTCTACTTCCACGAT | 22 | 61.53 | 54.54 | 7785 | 7763 |
| SLEV_30_LEFT | GAAAGCGTGGAGGAGGAAAAGG | 22 | 61.45 | 54.54 | 7662 | 7684 |
| SLEV_30_RIGHT | GGAACACGTCCACTCCACTTTT | 22 | 61 | 50 | 8070 | 8048 |
| SLEV_31_LEFT | CGCAACCCTGAAGCATGTTCAA | 22 | 62.21 | 50 | 7942 | 7964 |
| SLEV_31_RIGHT | GCCCCACTAACCCAGTACATCT | 22 | 61.49 | 54.54 | 8344 | 8322 |
| SLEV_32_LEFT | GTCCATACACGCCCAAAATCA | 23 | 61.67 | 47.82 | 8221 | 8244 |
| SLEV_32_RIGHT | TCACCATCGAGCTAGCTGATCC | 22 | 61.65 | 54.54 | 8640 | 8618 |
| SLEV_33_LEFT | AGTTGGGGAAAGGATACGGAGA | 22 | 60.56 | 50 | 8503 | 8525 |
| SLEV_33_RIGHT | TGGCTGTTCACCTTTGCTTTGA | 22 | 61.21 | 45.45 | 8893 | 8871 |
| SLEV_34_LEFT | ACCACTAGGAGTCGCCCAAATC | 22 | 62.06 | 54.54 | 8773 | 8795 |
| SLEV_34_RIGHT | CAAACTCCAAGAACCGAGCTCC | 22 | 61.43 | 54.54 | 9135 | 9113 |
| SLEVPrimers_38_LEFT | GATGGGAAAGCGCGAGAAGAAG | 22 | 61.86 | 54.54 | 8970 | 8992 |
| SLEVPrimers_38_RIGHT | ACCTTGTGGCGATAGGTCAGAT | 22 | 61.21 | 50 | 9339 | 9317 |
| SLEV_36_LEFT | TCCAGGAGGAAAGATGTACGCA | 22 | 61.14 | 50 | 9250 | 9272 |
| SLEV_36_RIGHT | TCAGGTCCATTCTTCCTCAGCC | 22 | 61.75 | 54.54 | 9643 | 9621 |
| SLEV_37_LEFT | CAACCTGGCCGTTCAACTGATA | 22 | 60.86 | 50 | 9517 | 9539 |
| SLEV_37_RIGHT | AGCTCATCCTGGCTCCTACATG | 22 | 61.60 | 54.54 | 9886 | 9864 |
| SLEV_38_LEFT | AGGACATTCAGGAGTGGAAACCT | 23 | 61.28 | 47.82 | 9750 | 9773 |
| SLEV_38_RIGHT | CTTCAATCCACACACGGTTCCA | 22 | 61.07 | 50 | 10143 | 10121 |
| SLEV_39_LEFT | TCAGCTGTCCCAGTCAACTG | 22 | 61.38 | 54.54 | 10023 | 10045 |
| SLEV_39_RIGHT | GCACTCCTCCTACCACATGAGT | 22 | 61.47 | 54.54 | 10386 | 10364 |
| SLEV_40_LEFT | CCACCTGGGCTGAGAACATCTA | 22 | 61.47 | 54.54 | 10254 | 10276 |
| SLEV_40_RIGHT | ACAGACAGCACCTTTAGCATGC | 22 | 61.45 | 50 | 10635 | 10613 |
| SLEV_41_LEFT | CAGGTAAACGGTGCTGTCTG | 22 | 61.43 | 54.54 | 10458 | 10480 |
| SLEV_41_RIGHT | CTGGTGTTGAAAAAGCAGGGGA | 22 | 61.28 | 50 | 10884 | 10862 |
